# Late Quaternary habitat suitability models for chimpanzees (*Pan troglodytes*) since the Last Interglacial (120,000 BP)

**DOI:** 10.1101/2020.05.15.066662

**Authors:** Christopher D. Barratt, Jack D. Lester, Paolo Gratton, Renske E. Onstein, Ammie K. Kalan, Maureen S. McCarthy, Gaëlle Bocksberger, Lauren C. White, Linda Vigilant, Paula Dieguez, Barrie Abdulai, Thierry Aebischer, Anthony Agbor, Alfred Kwabena Assumang, Emma Bailey, Mattia Bessone, Bartelijntje Buys, Joana Silva Carvalho, Rebecca Chancellor, Heather Cohen, Emmanuel Danquah, Tobias Deschner, Zacharie Nzooh Dongmo, Osiris A. Doumbé, Jef Dupain, Chris S. Duvall, Manasseh Eno-Nku, Gilles Etoga, Anh Galat-Luong, Rosa Garriga, Sylvain Gatti, Andrea Ghiurghi, Annemarie Goedmakers, Anne-Céline Granjon, Dismas Hakizimana, Nadia Haydar, Josephine Head, Daniela Hedwig, Ilka Herbinger, Veerle Hermans, Sorrel Jones, Jessica Junker, Parag Kadam, Mohamed Kambi, Ivonne Kienast, Célestin Yao Kouakou, Kouamé Paul N’Goran, Kevin E. Langergraber, Juan Lapuente, Anne Laudisoit, Kevin C. Lee, Fiona Maisels, Deborah Moore, Bethan Morgan, David Morgan, Emily Neil, Sonia Nicholl, Louis Nkembi, Anne Ntongho, Christopher Orbell, Lucy Jayne Ormsby, Liliana Pacheco, Alex K. Piel, Lilian Pintea, Andrew J. Plumptre, Aaron Rundus, Crickette Sanz, Volker Sommer, Tenekwetche Sop, Fiona A. Stewart, Jacqueline Sunderland-Groves, Nikki Tagg, Angelique Todd, Els Ton, Joost van Schijndel, Hilde VanLeeuwe, Elleni Vendras, Adam Welsh, José Francisco Carminatti Wenceslau, Erin G. Wessling, Jacob Willie, Roman M. Wittig, Nakashima Yoshihiro, Yisa Ginath Yuh, Kyle Yurkiw, Christophe Boesch, Mimi Arandjelovic, Hjalmar Kühl

## Abstract

**Aim:** Paleoclimate reconstructions have enhanced our understanding of how past climates may have shaped present-day biodiversity. We hypothesize that habitat stability in historical Afrotropical refugia played a major role in the habitat suitability and persistence of chimpanzees (*Pan troglodytes*) during the late Quaternary. We aimed to build a dynamic model of changing habitat suitability for chimpanzees at fine spatio-temporal scales to provide a new resource for understanding their ecology, behaviour and evolution.

**Location:** Afrotropics.

**Taxon:** Chimpanzee (*Pan troglodytes*), including all four subspecies (*P. t. verus, P. t. ellioti, P. t. troglodytes, P. t. schweinfurthii*).

**Methods:** We used downscaled bioclimatic variables representing monthly temperature and precipitation estimates, historical human population density data and an extensive database of georeferenced presence points to infer chimpanzee habitat suitability at 62 paleoclimatic time periods across the Afrotropics based on ensemble species distribution models. We mapped habitat stability over time using an approach that accounts for dispersal between time periods, and compared our modelled stability estimates to existing knowledge of Afrotropical refugia. Our models cover a spatial resolution of 0.0467 degrees (approximately 5.19 km^2^ grid cells) and a temporal resolution of every 1,000–4,000 years dating back to the Last Interglacial (120,000 BP).

**Results:** Our results show high habitat stability concordant with known historical forest refugia across Africa, but suggest that their extents are underestimated for chimpanzees. We provide the first fine-grained dynamic map of historical chimpanzee habitat suitability since the Last Interglacial which is suspected to have influenced a number of ecological-evolutionary processes, such as the emergence of complex patterns of behavioural and genetic diversity.

**Main Conclusions:** We provide a novel resource that can be used to reveal spatio-temporally explicit insights into the role of refugia in determining chimpanzee behavioural, ecological and genetic diversity. This methodology can be applied to other taxonomic groups and geographic areas where sufficient data are available.

## Introduction

Paleoclimate reconstructions have greatly enhanced our understanding of how biodiversity has been shaped globally since the late Quaternary, including inferences on range shifts, extinctions and the evolution of new lineages (Hewitt, 2000; 2004; Sandel et al., 2011; Svenning, Eiserhardt, Normand, Ordonez & Sandel, 2015). For example, climatic variability through time has played a major role in determining the uneven distribution of biodiversity across the Afrotropics (Demenocal, 1995; Maslin et al., 2014; Sepulchre et al., 2006; Trauth, Maslin, Deino & Strecker, 2005). Although a high proportion of African biodiversity is concentrated in a handful of forested hotspots and centres of endemism (Kingdon, 1990; Myers, Mittermeier, Mittermeier, Fonseca & Kent, 2000), these areas are geographically diffuse and historically complex, with idiosyncratic characteristics including heterogeneous topography, hydrological features, and highly dynamic and unique forest histories. Many Afrotropical forest lineages across different taxonomic groups are assumed to have tracked available habitat as it shifted throughout the Quaternary, contracting into refugia during glaciation, and often expanding during post-glacial periods to colonise (or re-colonise) new geographic areas. However, spatio-temporal reconstructions of Pleistocene forest refugia across the Afrotropics are often limited to broad scale maps without detailed information at local scales (Maley, 1996; Mayr & O’Hara, 1986). A more detailed quantification of the spatio-temporal distribution of forest refugia has been hampered by the lack of high-resolution paleoecological data with most previous reconstructions being limited to coarse spatial grains (Singarayer & Valdes, 2010) or to only a handful of temporal snapshots during the Quaternary (Hijmans, Cameron, Parra, Jones, & Jarvis, 2005). Here, we address this limitation by modelling the historical habitat suitability of chimpanzees (*Pan troglodytes*) across the Afrotropics using a comprehensive database of georeferenced presence points spanning their entire range (Fig. 1) and downscaled paleoclimate reconstructions since the Last Interglacial (120,000 BP).

**Fig. 1.**
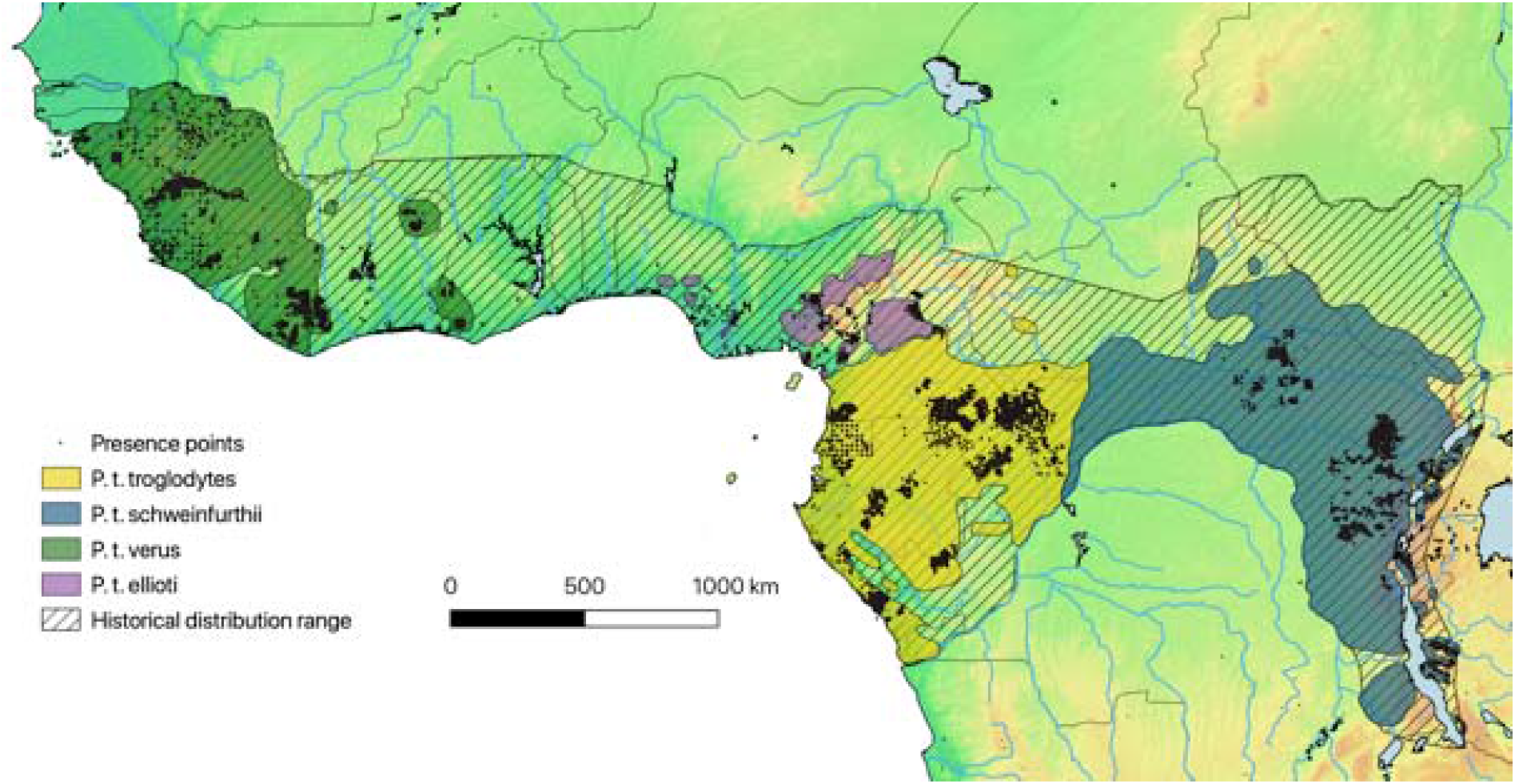
Current chimpanzee distribution and presence points extracted from the IUCN SSC A.P.E.S. database. Currently recognised subspecies ranges are depicted by coloured polygons (Humle et al., 2018), with black dots representing sampling localities (n=139,902 georeferenced presence points) used for species distribution modelling, historical distribution range (McBrearty & Jablonski, 2005) represented by diagonal shading. Country borders, topography (lowlands = green, highlands = brown) and major hydrological features are also shown.

Chimpanzees are an appropriate focal species for our approach due to their wide geographical distribution and high detectability. This means that their presence in an area is unlikely to be overlooked even in places which are only sporadically surveyed, as is the case for many poorly known and remote field sites across the Afrotropics. Their wide distribution encompasses high levels of ecological, behavioural, and genetic diversity amongst populations, providing a model system to study the availability of climatic refugia over time across diverse habitats and geographic regions chimpanzees occur across 21 African countries, in a wide variety of ecological conditions (from dense moist forest to arid savannah, from sea level up to an altitude of 2,800m) and varying levels of anthropogenic pressure (Humle, Maisels, Oates, Plumptre & Williamson, 2018). Four subspecies are currently described: *P. t. verus* from West Africa, *P. t. ellioti* from Nigeria and Cameroon, *P. t. troglodytes* from Central Africa and *P. t. schweinfurthii* from Central and Eastern Africa. Chimpanzee diversity is also apparent in their social organisation (community size ranges from 12 to 200, Langergraber et al. 2017), feeding ecology and diet (proportions and species of vertebrates, insects and fruits consumed varies among populations (Basabose, 2002; McGrew, Baldwin & Tutin, 1988; Nishida & Uehara, 1983; Wrangham, 1977), and range sizes (from around 3 km^2^ to over 60 km^2^ (Herbinger, Boesch, & Rothe, 2001; Pruetz & Bertolani, 2009; Wessling et al., 2018). Behavioural complexity in chimpanzees is highly variable, with diverse repertoires of tool use (e.g. sticks, stones and leaves to access insects, honey, meat, seeds and algae, water filtering) evident across forest and savannah populations (Galat-Luong, Galat & Nizinski, 2009; Kühl et al., 2019; Whiten et al., 1999). Additionally, genetic differentiation is also profound across the range, with four currently recognised subspecies and markedly variable population substructure and genetic diversity within each of these (CD Barratt, C Fontsere, JD Lester, pers. comm., Fünfstück et al., 2015; De Manuel et al., 2016; Mitchell et al., 2015; Prado-Martinez et al., 2013).

It is suspected that historical climatic changes and associated population size changes and migrations have been a major driver of chimpanzee genetic and cultural differentiation since at least the Last Interglacial (Prado-Martinez et al., 2013). To date, our understanding of the spatiotemporal climatic changes which could have influenced the mechanisms driving chimpanzee ecological, behavioural, and genetic diversity have been limited by the absence of information about historical climate change at fine spatial scales across the Afrotropics, a shortfall which this study aims to address. Specifically, we hypothesize that previously identified forest refugia played a major role in the habitat suitability and persistence of chimpanzees (*Pan troglodytes*) during late Quaternary climate fluctuations. Our results lay the foundations for a deeper understanding of eco-evolutionary processes responsible for chimpanzee distribution, diversification and resilience to climate change, providing a novel resource that can help to build testable hypotheses in a framework that can be broadly applied across taxonomic groups within Afrotropical forest ecosystems.

## Materials and Methods

### Reconstructing African paleoclimatic conditions back to the Last Interglacial

Paleoclimate conditions in Africa were obtained from Bell et al., (2017). These data reconstructed climate conditions since the Last Interglacial (120,000 BP) by downscaling the climate data at 1.25 degrees from Singarayer and Valdes (2010). These data are based on the Hadley Centre Coupled Model version 3 general circulation model (HadCM3), and represent 62 snapshots of global climatic conditions at 1,000 year intervals from the present back to the Last Glacial Maximum (LGM, ~22,000 BP), at 2,000 year intervals from the LGM to 80,000 BP, and at 4,000 year intervals from 80,000 BP back to the Last Interglacial. Monthly temperature and precipitation anomalies were downscaled to 0.2 degrees (~22.23 km) using a bilinear spline, and then to 0.0467 degrees (~5.19 km) globally using a bicubic spline. Eight climate variables were reconstructed: mean annual temperature (bioclim_01), temperature seasonality (bioclim_04), mean temperature of the warmest (bioclim_10) and coldest quarters (bioclim_11), mean annual precipitation (bioclim_12), precipitation seasonality (bioclim_15), and precipitation of the wettest (bioclim_16) and driest quarters (bioclim_17) for each snapshot. We restricted our geographic extent to encompass both the contemporary and potential historical habitat suitability of chimpanzees (McBrearty and Jablonski, 2005), from −18 to 32 degrees longitude and −10 to 16 degrees latitude. This geographic extent includes the current distribution range of bonobos (*Pan paniscus*), the sister species of chimpanzees, spatially separated from the latter by the Congo River. We included the bonobo distribution in our models due to previous detection of historic admixture between chimpanzees and bonobos based on genomic data (De Manuel et al., 2016; Kuhlwilm, Han, Sousa, Excoffier, & Marques-Bonet, 2019), suggesting that chimpanzees may have occasionally been found in these areas. Spatial downscaling analyses were conducted with the ‘climates’ R package (VanDerWal, Beaumont, Zimmerman & Lorch, 2011) in R (R Core Team, 2019).

To account for anthropogenic effects on chimpanzee habitat suitability, we also included a spatial layer available from the HyDE database (Klein Goldewijk, Beusen, Van Drecht, & De Vos, 2011), which provides information on modelled historical human population density based on population and agricultural data. We extended the human population densities back in time from 12,000 BP (i.e., the last available data from the HyDe database) uniformly to the Last Interglacial (120,000 BP) and downscaled it so that it matched the temporal and spatial scale of our paleoclimate reconstructions. We selected these climatic and anthropogenic variables to reflect biologically informative conditions likely to have influenced chimpanzee habitat suitability, and because these variables were available at paleoclimatic timescales.

### Species distribution modelling of chimpanzees

We compiled a database of 139,902 georeferenced chimpanzee presence point records (nests, sightings, faeces, footprints) from our own fieldwork, from the IUCN SSC A.P.E.S. database, a collaborative initiative to centralize great ape population surveillance data (http://apes.eva.mpg.de/) and from the Pan African Programme: The Cultured Chimpanzee (http://panafrican.eva.mpg.de/). In total this data represents over fifty collaborating wildlife research institutions across the entire distributional range of chimpanzees. After checking the collinearity of our climatic and anthropogenic variables based on contemporary conditions, we reduced our nine variables to six due to high co-correlations (Pearson’s r > 0.6). Our retained variables were: annual mean temperature, temperature seasonality, annual precipitation, precipitation of wettest quarter, precipitation of driest quarter, and human density. We then constructed an ensemble species distribution model (SDM) representing the contemporary habitat suitability of chimpanzees (i.e. all combined subspecies) using the “sdm” R package version 1.0-46 (Naimi & Araújo, 2016).

We followed recommended guidelines for constructing SDMs (Merow, Smith and Silander, 2013, Merow et al., 2014, Araujo, 2019), first rarefying the presence data so that no points were within 10 km of one another to reduce spatial autocorrelation, resulting in 1,677 unique presence points. Background points (sometimes referred to as ‘pseudoabsences’ in the SDM literature) were generated randomly from within a 0.5 degree buffer radius of presence points to avoid sampling areas where no surveys of chimpanzees have been conducted (Fig. S1), with the number of points equal to the number of presence points in our SDMs. We selected background points in this way to emphasize factors locally relevant in distinguishing suitable from unsuitable habitat, while adequately sampling the range of climatic conditions for our study species (VanDerWal, Shoo, Graham, & Williams, 2009). Rather than relying on individual modelling algorithm approaches, we built ensemble models combining multiple replicates of several different modelling algorithms to represent alternate possible states of the system being modelled (Araújo & New, 2007). Due to their combined power, ensemble models are widely accepted to provide more accurate results than single models (Forester, Dechaine, & Bunn, 2013). We used five cross-validated replicates each of 14 different modelling algorithms available in the sdm R package (Bioclim, Bioclim.dismo, Boosted Regression Trees, Classification and Regression Trees, Flexible Discriminant Analysis, Generalized Additive Model, Generalized Linear Model, Multivariate Adaptive Regression Spline, Maximum Entropy, Maximum Entropy-like, Mixture Discriminant Analysis, Random Forest, Recursive Partitioning and Regression Trees, Support Vector Machine), partitioning data into training (70%) and testing (30%). We evaluated the performance of each of the five replicates of the 14 modelling algorithms performance based on the Area Under the Curve of a Receiver Operating Characteristics plot (AUC, Fielding & Bell, 1997) and True Skill Statistic (TSS, Allouche, Tsoar, & Kadmon, 2006). Modelling algorithms that performed adequately (AUC > 0.8, and TSS > 0.5, Bell et al., 2017) were then combined in an ensemble model prediction using five replicates of each algorithm, weighting the contribution of each modelling algorithm to the ensemble model by their AUC. For each modelling algorithm and the ensemble model we also measured the permutation importance of predictor variables across each model iteration using the “getVarImp” function in the ‘sdm’ R package.

### Paleoclimatic niche models and habitat stability

To identify geographic areas where high habitat suitability for chimpanzees has remained stable since the Last Interglacial (i.e., to identify refugia where the effects of climate change have been less pronounced than surrounding areas), we projected our SDM back in time so that we had a habitat suitability estimate for each pixel at each paleoclimate snapshot. We projected our contemporary ensemble model onto the 62 snapshots of downscaled paleoclimate reconstructions and human density data since the Last Interglacial using the “predict” function of the ‘sdm’ R package. For consistency, we assumed that the fundamental ecological niche of chimpanzees represented by our predictor variables remained relatively constant over this period, which we believe is a reasonable assumption given the large geographical extent and wide range of climatic conditions that our spatial and climatic data accounts for, though we acknowledge there may be exceptions at local scales over such long temporal intervals where fundamental niches have shifted over time. Based on these per pixel estimates of habitat suitability over time, we then evaluated suitability changes through space and time, using two approaches: Static stability (Hugall, Moritz, Moussalli, & Stanisic, 2002), which does not consider dispersal between pixels, and Dynamic stability (Barratt et al., 2017; Graham, VanDerWal, Phillips, Moritz, & Williams, 2010; Rosauer, Catullo, Vanderwal, & Moussalli, 2015), which uses a graph theoretic approach with a defined dispersal rate to model how chimpanzees may track suitable climate conditions across time. Using the dynamic stability approach, a pixel is considered as stable as long as pixels within the defined dispersal distance have suitable climate in adjacent time steps. For the static stability estimate, we summed the negative log of suitability through time for each pixel, and for the dynamic stability estimate we allowed a conservative five metres/year dispersal rate, which amounts to a maximum dispersal distance of 600 km over the 120,000 year time period for chimpanzee populations since the Last Interglacial, approximately matching the historical distribution inferred by McBrearty & Jablonski (2005) shown in Fig. 1. All static and dynamic stability surfaces were calculated using modified R scripts from Graham et al. (2010) provided by Jeremy VanDerWal. We then visually compared our modelled areas of long-term habitat stability (i.e. refugia) to those previously inferred in the literature (Huntley, Keith, Castellano, Musher & Voelker, 2019; Maley, 1996; Mayr & O’Hara, 1986).

### Sensitivity analyses

Our final contemporary chimpanzee habitat suitability model (Fig. 2) represents all four currently recognised subspecies combined (Humle et al., 2018), but we also repeated the SDM, paleoclimate projections and stability estimations described above for each subspecies separately to ensure that our model shown in Fig. 1 did not underpredict habitat suitability at local scales that may be ecologically important for each subspecies. The number of presence points (and background points) used for the subspecies, *P. t. verus, P. t. ellioti, P. t. troglodytes* and *P. t. schweinfurthii* were 519, 134, 663, and 451, respectively). As a further sensitivity analysis we also repeated all analyses (i.e., combined subspecies together and for each subspecies separately) using presence data spatially rarefied so that no points were within 25 km of each other to assess whether spatial bias in the presence data was influencing the output models (number of presence points and background points for combined subspecies and for *P. t. verus, P. t. ellioti, P. t. troglodytes* and *P. t. schweinfurthii* separately were 658, 225, 57, 212 and 164, respectively) (Fig. S1). To statistically test how well our contemporary habitat suitability models and stability models (dynamic and static stability) matched one another across sensitivity analyses (with different spatial rarefaction in the input presence points) we conducted Pearson’s correlation tests in R using the ‘raster’ package version 3.1-5 (Hijmans et al., 2011). We also visually compared our model outputs against other published contemporary habitat suitability estimates available for chimpanzees which are available at different spatial resolutions and extents than our models (Heinicke et al., 2019; Jantz, Pintea, Nackoney, & Hansen, 2016; Junker et al., 2012; Sesink Clee et al., 2015; Strindberg et al., 2018).

**Fig. 2.**
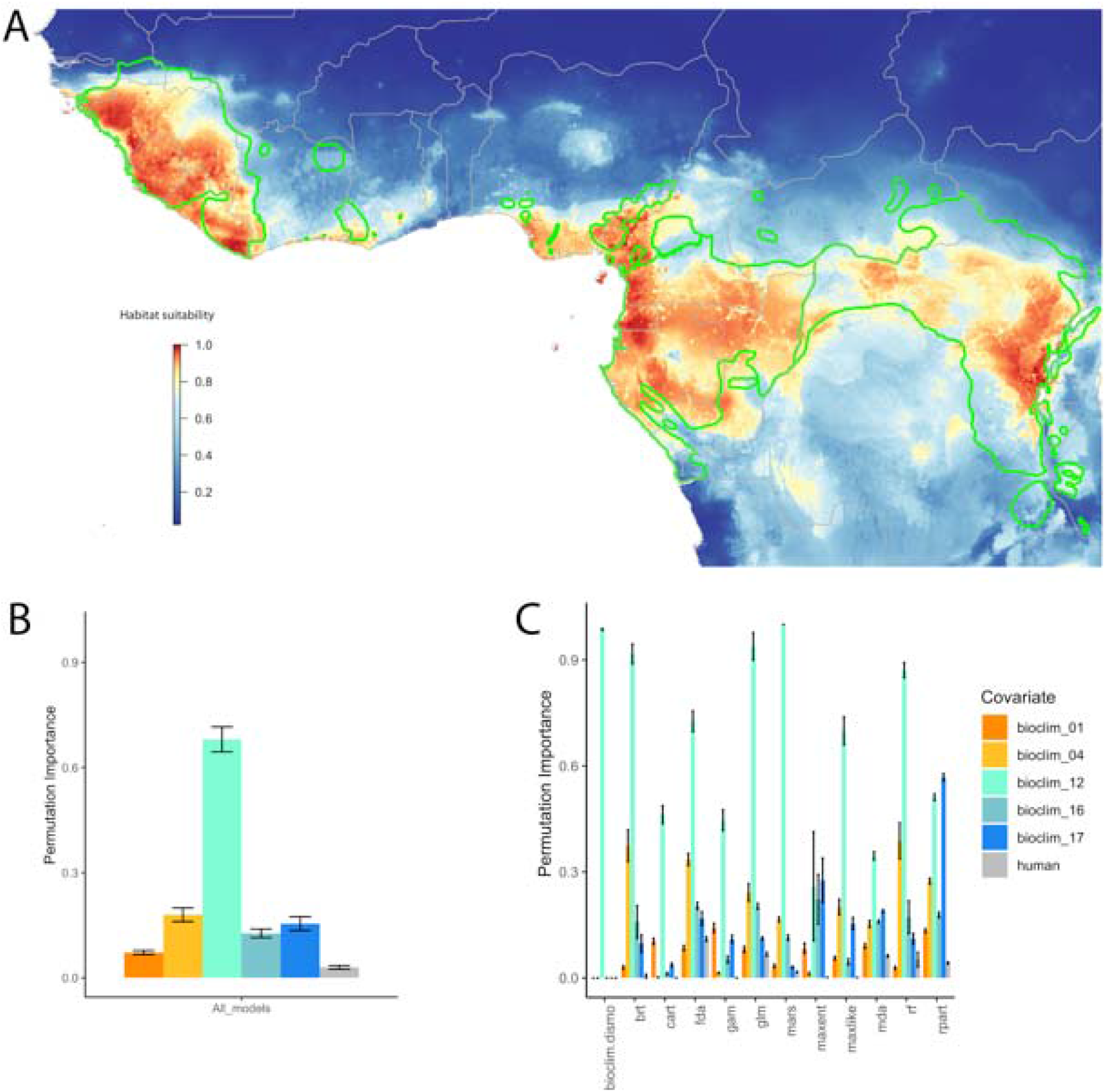
**A)** Contemporary habitat suitability model for chimpanzees (ensemble model of 5 crossvalidated replicates of 12 modelling algorithms) and predictor variable permutation importance B) averaged across all modelling algorithms, and C) per individual modelling algorithm. Axes represents predictor variable permutation importance (y axis), for each modelling algorithm (x axis), bioclim_01 = Mean annual temperature, bioclim_04 = temperature seasonality, bioclim_12 = mean annual precipitation, bioclim_16 = precipitation of the wettest quarter, bioclim_17 = precipitation of the driest quarter, human = human density. Country borders (grey lines) and chimpanzee subspecies ranges (green lines, Humle et al., 2018) are also shown.

## Results

### Contemporary chimpanzee habitat suitability

Our contemporary habitat suitability model performed generally well at species level (i.e. all four subspecies combined), with AUC > 0.8 and TSS > 0.5 for 12 of the 14 tested modelling algorithms (Table 1). This was similar for sensitivity analyses on subspecies models *P. t. schweinfurthii* (*n*=13), *P. t. troglodytes* (*n*=13) and *P. t. verus* (*n*=11) (Table S1). Bioclim and bioclim.dismo algorithms performed poorly for most models (i.e. low AUC < 0.8, TSS < 0.5) and were therefore excluded from the ensemble models. For *P. t. elliotti*, all 14 modelling algorithms passed our AUC and TSS thresholds and so were retained in the ensemble model for this subspecies. The best performing modelling algorithms (i.e., higher AUC and TSS) were most often Maxent, Generalized Additive Models, and Random Forest (Table 1, Table S1). These results were also consistent when performing the sensitivity analyses using only presence points rarefied to a minimum distance of 25 km, indicating that our results are robust against spatial bias in the input data (Table S1), albeit with generally lower AUC and TSS, most probably due to the reduced number of presence points used to train the models.

**Table 1.** Performance of each individual modelling algorithm based on five cross-validated replicates for full species (*P. troglodytes*) and each of the four currently recognised subspecies using presence points rarefied to minimum 10 km distance from one another. Numbers indicate Area Under the Curve of a Receiver Operating Characteristics plot (AUC) and True Skill Statistic (TSS) for each modelling algorithm, with high values indicating better model performance. ***Bold italic*** values represent AUC < 0.8 or TSS <0.5, the minimum threshold we set for that modelling algorithm to be included in the ensemble models. Modelling algorithm abbreviations – bioclim: Bioclim, bioclim.dismo: Bioclim from the dismo R package, brt: Boosted Regression Trees, cart: Classification and Regression Trees, fda: Flexible Discriminant Analysis, gam: Generalized Additive Model, glm, Generalized Linear Model, mars: Multivariate Adaptive Regression Spline, maxent: Maximum Entropy, maxlike: Maximum Entropy-like, mda: Mixture Discriminant Analysis, rf: Random Forest, rpart: Recursive Partitioning and Regression Trees, svm, Support Vector Machine.

| Modelling algorithm | *P. troglodytes* |  | *P. t. verus* |  | *P. t. ellioti* |  | *P. t. troglodytes* |  | *P. t. schweinfurthii* |  |
| --- | --- | --- | --- | --- | --- | --- | --- | --- | --- | --- |
| evaluation metric | AUC | TSS | AUC | TSS | AUC | TSS | AUC | TSS | AUC | TSS |
| bioclim | ***0.63*** | ***0.23*** | ***0.64*** | ***0.3*** | ***0.8*** | 0.59 | ***0.75*** | ***0.45*** | ***0.73*** | ***0.41*** |
| bioclim.dismo | ***0.72*** | ***0.34*** | ***0.76*** | ***0.46*** | ***0.94*** | 0.86 | ***0.89*** | 0.73 | ***0.85*** | 0.6 |
| brt | 0.8 | 0.51 | 0.84 | 0.57 | 0.97 | 0.88 | 0.9 | 0.68 | 0.89 | 0.61 |
| cart | 0.84 | 0.59 | 0.86 | 0.62 | 0.92 | 0.87 | 0.94 | 0.83 | 0.89 | 0.72 |
| fda | 0.81 | 0.51 | 0.83 | 0.58 | 0.97 | 0.85 | 0.85 | 0.67 | 0.89 | 0.64 |
| gam | 0.87 | 0.62 | 0.88 | 0.65 | 0.98 | 0.94 | 0.97 | 0.88 | 0.92 | 0.73 |
| glm | 0.82 | 0.56 | 0.82 | 0.57 | 0.97 | 0.88 | 0.88 | 0.71 | 0.91 | 0.67 |
| mars | 0.87 | 0.63 | 0.88 | 0.65 | 0.98 | 0.93 | 0.95 | 0.84 | 0.92 | 0.72 |
| maxent | 0.87 | 0.61 | 0.89 | 0.64 | 0.98 | 0.94 | 0.95 | 0.83 | 0.93 | 0.76 |
| maxlike | 0.82 | 0.57 | 0.84 | 0.59 | 0.97 | 0.89 | 0.86 | 0.73 | 0.88 | 0.65 |
| mda | 0.83 | 0.54 | 0.84 | 0.61 | 0.97 | 0.91 | 0.91 | 0.75 | 0.88 | 0.65 |
| rf | 0.94 | 0.77 | 0.9 | 0.74 | 0.98 | 0.95 | 0.98 | 0.9 | 0.97 | 0.89 |
| rpart | 0.85 | 0.63 | 0.86 | 0.62 | 0.95 | 0.9 | 0.95 | 0.84 | 0.89 | 0.74 |
| svm | 0.91 | 0.68 | ***0.79*** | 0.66 | 0.97 | 0.9 | 0.97 | 0.86 | 0.93 | 0.73 |
| Total n models in ensemble | 12 |  | 11 |  | 14 |  | 13 |  | 13 |  |

The contemporary habitat suitability model for the full species (Fig. 2A) showed more extensive areas of high suitability compared to sensitivity analyses when subspecies were modelled separately, and suitability was also lower using the 25 km rarefied input presence points (Fig. S2, Table S2). Our contemporary habitat suitability models for chimpanzees (Fig. 2A) approximately matched those previously published (Heinicke et al., 2019; Jantz et al., 2016; Junker et al., 2012; Sesink Clee et al., 2015; Strindberg et al., 2018) but there were several differences, likely due to the different variables used to build models that are unavailable or inappropriate at the timescales we projected (e.g., forest cover and change, distance to roads/rivers, disease dynamics), and the different spatial resolution and extent of the models. For example, our models predicted much higher habitat suitability in west Africa for *P. t. verus* across parts of Sierra Leone, Liberia and Guinea than some previous work (Jantz et al., 2016; Junker et al., 2012), though they are similar to the modelled chimpanzee density patterns reported by Heinicke et al. (2019). Our models tended to underpredict suitability for *P. t. troglodytes* in a small part of central Gabon but generally matching the remainder of this subspecies range compared to Strindberg et al. (2018). For *P. t. ellioti*, modelled habitat suitability mirrors the predicted chimpanzee distributions shown by Sessink-Clee et al. (2015) very closely, though the areas of higher habitat suitability are slightly larger in our models. For *P. t. schweinfurthii*, our models are concordant with those of Junker et al. (2012) and Jantz et al. (2016).

Summarizing variable permutation importance indicated that precipitation-related predictors (bioclim_16 – precipitation of wettest quarter, bioclim_17 – precipitation of driest quarter, and in particular bioclim_12 – annual rainfall) were more important than temperature-related predictors (bioclim_01 – annual mean temperature, bioclim_04 – temperature seasonality) and human population density in explaining chimpanzee habitat suitability (Fig. 2B, C). These results were consistent across all sensitivity analyses including models using the 25 km rarefied input presence points and for separate subspecies models except for *P. t. ellioti* (bioclim_16 with the highest permutation importance rather than bioclim_12) and *P. t. schweinfurthii*, which had similar variable importance between all temperature and precipitation predictor variables, and low importance for human density (Fig. S3).

### Habitat suitability through time and identifying potential refugia

Visual inspection of the dynamic stability estimates (Fig. 3) indicated that the areas that have remained consistently more climatically stable closely match the forest refugia posited by previous studies (Maley, 1996; Mayr & O’Hara, 1986), with our data providing estimates at finer spatial scales than previous work. Dynamic stability estimates that accounted for dispersal showed larger areas of suitability than static stability estimates (Fig. S4, S5, Table 2), though both highlighted the same general regions as areas of long-term habitat stability. Our stability estimates corroborated that refugia suggested in previous studies have remained more stable than surrounding areas; all three refugia in the Upper Guinea Forests, both refugia in the Cameroon Volcanic Line and Lower Guinea Forests, two of three refugia through Gabon and Congo-Brazzaville, the refuge within the bonobo distribution in central DRC, and several parts of the fragmented refugia across the Albertine Rift (Maley 1996; Mayr & O’Hara, 1986), however missing some of the central African microrefugia described by Leal (2004). Our results suggest some of these refugia may have been previously underestimated in size, with areas throughout the Upper Guinea Forests in Sierra Leone (Loma Mountains), Liberia (Kpo Mountains) and southern Guinea (Massif du Ziama and Nimba Mountains), areas surrounding the mountains of the Cameroon Volcanic Line (Mt. Cameroon, Manengouba, Bamboutos, Lebialem Highlands), the Mts. De Cristal - Massif du Chaillu mountain chain that runs from northwest to southeast from central Equatorial Guinea through northern and Central Gabon to the border with the Republic of Congo, and parts of the Albertine Rift and surrounding forests (Kahuzi-Biega, Ituri, Bwindi, Ruwenzori Mountains, Kibale) in particular that lie outside previously estimated refugia. Habitat suitability and stability (dynamic and static) estimates across our sensitivity analyses using different levels of presence points spatial rarefaction, 10 km vs. 25 km) were also highly consistent (Pearson’s r > 0.971 in all cases, Table S2, Fig. S4, S5). We provide an animated image of habitat suitability over time for each of the 62 paleoclimatic snapshots for the full species for visualization (Fig. S6).

**Fig. 3.**
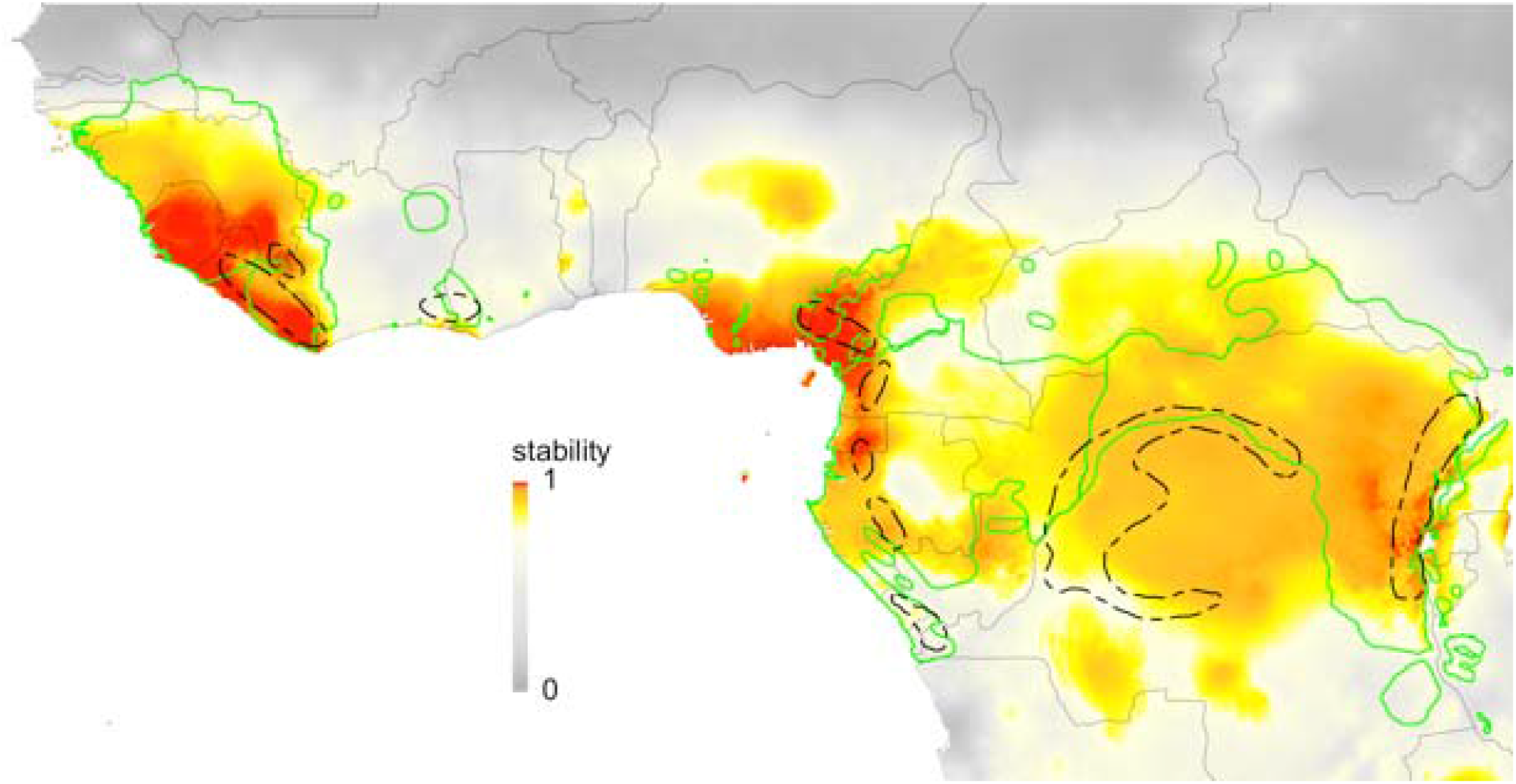
Stability of chimpanzee habitat suitability (0= low suitability stability to 1=high suitability stability, using the full species dataset) over 62 snapshots of paleoclimate reconstructions representing the past 120,000 years using the Dynamic stability approach (dispersal rate of 5 m/year, amounting to a total possible dispersal of 600 km over the 120,000 years) compared to previous estimates of forest refugia (dotted black lines, Maley, 1996). Country borders (grey lines) and chimpanzee subspecies ranges (green lines, Humle et al., 2018) are also shown.

## Discussion

We hypothesized that Pleistocene forest refugia played a major role in the persistence of chimpanzees (*Pan troglodytes*) during late Quaternary climate fluctuations, testing this by building spatio-temporally explicit models of chimpanzee habitat suitability across Africa over the last 120,000 years, since the Last Interglacial. We found several areas of high habitat suitability that have remained stable over this time period, largely matching estimates of putative refugia from previous studies (Maley, 1996; Mayr & O’Hara, 1986). Our results suggest that for chimpanzees, most of these previous refugia are underestimated in size, implying that additional geographic areas have potentially supported suitable habitats during historical climate oscillations, corresponding to the estimated extent of their historical range (McBrearty & Jablonski, 2005). These extended refugial areas include parts of the Upper Guinea Forests in Sierra Leone (Loma Mountains), Liberia (Kpo Mountains) and southern Guinea (Massif du Ziama and Nimba mountains), and in and around three important African mountain chains: the Lower Guinea Forests surrounding the mountains of the Cameroon Volcanic Line, the Monts de Cristal-Massif du Chaillu mountain chain that passes through central Equatorial Guinea, northern and central Gabon, to the border with the Republic of Congo, and the Albertine Rift. Below we discuss how our results can complement current knowledge about refugia across the Afrotropics for chimpanzees, noting some caveats for their interpretation, and identify some future directions for how our novel resource may be integrated with other data types to build testable hypotheses about the complex contemporary patterns of ecological, behavioural, and genetic diversity exhibited by chimpanzee populations across their range, and for biodiversity in tropical regions more generally.

### Understanding historical and recent factors shaping contemporary biodiversity patterns

Many previous studies have shown that climate change throughout the late Quaternary had a profound influence on the contemporary distribution and structure of species and populations globally (Davis & Shaw, 2001; Ellegren & Galtier, 2016; Hewitt, 2000; Hewitt, 2004; Sandel et al., 2011; Svenning et al., 2015). In Africa it has been hypothesized that forest refugia, often in mountainous regions, are important areas for buffering the effects of climate change on biodiversity over paleoclimatic scales, supporting rare species and unique genetic variants (Fjeldsaå & Lovett, 1997; Mayr & O’Hara, 1986). Although initially these refugia were identified using species distributional patterns for birds and plants, an increasing number of recent studies have utilized molecular data to infer the geographical location of intraspecific diversity (i.e lineages or populations within species) across the Afrotropics. These studies have demonstrated contemporary phylogeographic patterns that suggest broadly concordant historical refugia in mammals (Bohoussou et al., 2015; Bryja, Mikula, Patzen, et al., 2014; Bryja, Mikula, Šumbera, et al., 2014; Bryja et al., 2017; Gaubert et al., 2016; Mizerovská et al., 2019; Nicolas et al., 2011), including other primates, (Anthony et al., 2014; Clifford et al., 2004; Gonder et al., 2011; Pozzi, 2016; Telfer et al., 2003), amphibians (Charles et al., 2018; Leaché et al., 2019; Portik et al., 2017), and plants (Faye et al., 2016; Hardy et al., 2013; Piñeiro, Dauby, Kaymak, & Hardy, 2017; Piñeiro et al., 2019), albeit with some differences between species due to different ecological characteristics and idiosyncratic responses to climatic changes (Lowe, Harris, Dormontt, & Dawson, 2010).

Our results are concordant with most of the broadly defined historical refugia for the Afrotropics (Maley, 1996; Mayr & O’Hara, 1986) but suggest that these areas are likely to be underestimated in their extent for chimpanzees. This may well be due to the ecological and behavioural flexibility exhibited by chimpanzees (Kühl et al., 2019; Wessling et al., 2018; Whiten et al., 1999), with their ability to tolerate drier savannah-like conditions, for example at the edges of their current distribution range where forest interdigitates with savannah, resulting in a broad ecological niche. This flexibility may also account for our estimates of chimpanzee habitat suitability and stability over time being generally higher when modelling all four subspecies combined compared to modelling each subspecies separately, as the ecological niche of each subspecies is narrower than that of the whole species. Since the turn of the 20^th^ century, the higher deforestation rates around the edges of chimpanzee range (e.g., in West Africa) than those at the core of their range (Aleman, Jarzyna, & Staver, 2018), may have caused the local extirpation of some populations, and potentially stimulated ecological and behavioural diversification in others. Historical climate changes combined with these more recent anthropogenic effects may help to explain some of the complex patterns of behavioural and cultural diversity described in chimpanzees across different parts of Africa (Kühl et al., 2019; Whiten et al., 1999) and particularly why some behaviours are geographically disjunct (e.g., accumulative stone throwing, Kühl et al. (2016), algae fishing (Boesch et al., 2017). However, these complex patterns of diversity are further complicated by multiple factors, including disease dynamics (Leroy et al., 2004), the bushmeat trade (Bennett et al., 2007), and local genetic adaptations (Schmidt, De Manuel, Marques-Bonet, Castellano, & Andrés, 2019).

### Caveats

Although our reconstructions of paleoclimatic refugia align with those of Maley (1996) and Mayr and Hara (1986), and with more recent molecular evidence in vertebrates and plants, we advocate caution in their interpretation despite the statistical robustness across our sensitivity analyses. Firstly, our approach uses a limited number of climatic and anthropogenic variables, and although we generally find similar contemporary habitat suitability predictions and/or density predictions as other studies (Heinicke et al., 2019; Jantz et al., 2016; Junker et al., 2012; Sesink Clee et al., 2015; Strindberg et al., 2018), differences exist, likely due to the discordance of the predictor variables used between ours and other approaches. Unfortunately, we lack sufficient spatio-temporal data on important predictor variables such as historical vegetation types, hydrological features and disease dynamics over the past 120,000 years, which may be important for predicting habitat suitability of chimpanzees at local scales. Due to this lack of data we were forced to restrict our predictor variables to those with the appropriate temporal resolution (i.e., paleoclimate and modelled historical human density). Secondly, it should be noted that the climate data we use are statistically downscaled from coarser resolution imagery from a single general circulation model (GCM) using locally available climate data (HadCM3). Though these types of data are widely used in a number of other studies (e.g., Bell et al., 2017; Graham et al., 2010; Rosauer et al., 2015), their accuracy may be improved by combining and testing different predictions in the ensemble model to capture variation between GCMs. Underlying assumptions regarding the HyDE human density data should also be acknowledged, mainly surrounding the interpolation techniques used to complete gaps in data collection (see https://themasites.pbl.nl/tridion/en/themasites/hyde/uncertainty/index-2.html). Thirdly, our SDMs use available chimpanzee presence points from across their entire range, but there remain significant sampling gaps affecting models because of a lack of exploration due to unstable political situations and disease outbreaks (e.g., parts of the *P. t. schweinfurthii* distribution range in DRC, Central African Republic and South Sudan). Finally, related to our discussion about ecological and behavioural flexibility we would like to emphasize that our models should not be interpreted as areas of forest or savannah stability per se, but rather as areas that are suitable for chimpanzees, defined by their ecological niche.

### Future directions for understanding diversification mechanisms

By modelling habitat suitability and stability over the last 120,000 years at high spatial and temporal resolution, we provide a novel resource to investigate the diversification mechanisms underlying patterns of ecological, behavioral and genetic diversity in chimpanzees. We suggest that future research should exploit our approach to generate testable hypotheses about chimpanzee population diversity by combining our spatio-temporally explicit models with ecological, behavioural, and molecular data. For example, habitat suitability fluctuations over time for a given set of populations may be explicitly tested against home range sizes, measures of genetic diversity or estimates of effective population size changes over time, or to the presence or absence of certain behaviours. Furthermore, hypotheses about genetic connectivity and behavioural transmission between populations across landscapes over time could be tested by integrating spatio-temporally explicit connectivity metrics based on circuit theory (Dickson et al., 2019; McGarigal & Marks, 1995; McRae & Beier, 2007) with empirical genetic and behavioural data (e.g. genetic differentiation F_ST_, G_ST_ and cultural F_ST_, Bell, Richerson, & McElreath, 2009). Connecting these concepts would also facilitate a predictive framework whereby the genetic, behavioural, and ecological characteristics of populations in poorly known geographic regions could be estimated, before validation with new empirical data as it becomes available.

Beyond correlative approaches such as those described above, mechanistic approaches would enable a deeper exploration of biogeographical patterns and processes that have affected chimpanzees, especially with the recent availability of comprehensive behavioural (Kühl et al., 2019) and molecular data (CD Barratt, C Fontsere, JD Lester, pers. comm.) for a large number of wild chimpanzee populations. Rapid developments in the generation and analysis of genomewide molecular data over the past decade have revealed detailed demographic histories, enabling the identification of diversification mechanisms due to forest refugia, which are characterised by divergence, isolation, and secondary contact as refugial habitats fragment and reconnect with each other during glacial cycles (Barratt et al. 2018; Charles et al., 2018; Leaché et al., 2019; Portik et al., 2017, Feng, Ruhsam, Wang, Li & Wang, in press). The ability to distinguish signals of forest refugia from other diversification mechanisms such as landscape barriers, ecological gradients, and anthropogenic habitat fragmentation, would represent a powerful approach for gaining a more mechanistic understanding of population diversification.

### Conclusions

Deeper insights into historical climate change in highly diverse tropical regions are essential to understand their contribution towards shaping current biodiversity patterns, and how biodiversity loss might be predicted given future projections of anthropogenic and climate change. By increasing the spatial resolution and number of time periods used to project SDMs, the novel resource we present here is able to improve on existing paleoclimate data that are temporally limited to build fine scale models of changing habitat suitability for species through time accounting for their dispersal ability. Integrating this resource with other data types in the future (e.g. ecological, behavioural and molecular) will help to increase our general understanding of how climate change impacts biodiversity, and how we may mitigate against predicted biodiversity loss in the future.

## Supporting information

Supplemental Information Figs 1-5, Tabs S1-S2

## Acknowledgements

We are indebted to several organisations including the Rwanda Development Board, Rwanda, Agence Nationale des Parcs Nationaux, Gabon, Instituto da Biodiversidade e das Áreas Protegidas (IBAP), Guinea□Bissau, Ministère de l’Enseignement Supérieur et de la Recherche Scientifique and the Office Ivoirien des Parcs et Réserves of Côte d’Ivoire, Ministère des Forêts et de La Faune, Cameroon, Institut Congolais pour la Conservation de la Nature (ICCN), Democratic Republic of the Congo, Ministère de la Recherche Scientifique, Democratic Republic of the Congo, Ministry of Agriculture, Forestry and Food Security, Sierra Leone, Forestry Development Authority of Liberia, Liberia and the Wildlife Conservation Society (WCS). We would also like to acknowledge numerous individuals for valuable input to this project: Ekwoge E. Abwe, Samuel Angedakin, Floris Aubert, Emmanuel Ayuk Ayimisin, Nsengiyumva Barakabuye, Donatienne Barubiyo, Barca Benjamin, Richard Bergl, Anna Binczik, Hedwige Boesch, Matthieu Bonnet, Terry Brncic, Damien Caillaud, Kenneth Cameron, Genevieve Campbell, Chloe Cipoletta, Katherine Corogenes, Charlotte Coupland, Pauwel De Wachter, Karsten Dierks, Emmanuel Dilambaka, Dervla Dowd, Andrew Dunn, Villard Ebot Egbe, Atanga Ekobo, J. Michael Fay, Joel Gamys, Jessica Ganas, Nicolas Granier, Liz Greengrass, John Hart, David Hebditch, Annika Hillers, Inaoyom Imong, Kathryn J. Jeffery, Mbangi Kambere, Noelle Kumpel, Deo Kujirakwinja, Vincent Lapeyre, Bradley Larson, Stephanie Latour, Vera Leinert, Paul Marchesi, Giovanna Maretti, Sergio Marrocoli, Rumen Martín, Amelia Meier, Nadia Mirghani, Felix Mulindahabi, Mizuki Murai, Stuart Nixon, Protais Niyigaba, Louis Nkembi, Emmanuelle Normand, Nicolas Ntare, Leonidas Nzigiyimpa, Robinson Orume, Leon Payne, Charles Petre, Kathryn Phillips, Jodie Preece, Frank Princee, Sebastien Regnaut, Alexis Kalinda Salumu, Emma Stokes, Alexander Tickle, Angelique Todd, Clement Tweh, Jeremy VanDerWal, Virginie Vergnes, Magloire Kambale Vyalengerera, Ymke Warren^†^, Klaus Zuberbühler. CDB, REO, HSK further acknowledge the support of the German Centre for Integrative Biodiversity Research (iDiv) Halle□Jena□Leipzig funded by the Deutsche Forschungsgemeinschaft (DFG, German Research Foundation)—FZT 118. This project has been partially conducted in the framework of the iDiv□Flexpool—the internal funding mechanism of iDiv. The scientific results have been computed at the High-Performance Computing (HPC) Cluster EVE, a joint effort of both the Helmholtz Centre for Environmental Research - UFZ (http://www.ufz.de/) and the German Centre for Integrative Biodiversity Research (iDiv) Halle-Jena-Leipzig (http://www.idiv-biodiversity.de/).

## Data availability

We provide our scripts and data for reproducibility as a DRYAD package. The package contains the raw data along with scripts to perform all analyses, build and run the models, and plot the results. The full distribution data for chimpanzees used in the analyses is available on request from the IUCN SSC A.P.E.S. database manager (http://apes.eva.mpg.de/).

## Biosketch

C.D.B. is a postdoctoral researcher at the German Centre for Integrative Biodiversity Research (iDiv) Halle-Jena-Leipzig, focused mainly on vertebrate biogeography, evolution and conservation. He is affiliated with the Department of Primatology within the Max Planck Institute for Evolutionary Anthropology as part of the Pan African Programme: The Cultured Chimpanzee. Author Contributions: C.D.B., R.E.O., J.L, P.G., C.B., M.A., and H.K conceived the ideas, all authors contributed presence point data, and J.V. contributed downscaled paleoclimate data for Africa. C.D.B analysed the data, and led the writing along with R.E.O and H.K, with contributions from all authors.

