## Supplemental Information Figs 1-5, Tabs S1-S2 for "Late Quaternary habitat suitability models for chimpanzees (*Pan troglodytes*) since the Last Interglacial (120,000 BP)"

**Short running title:** Habitat suitability of chimpanzees to the LIG

Christopher D. Barratt^1,2^, Jack D. Lester^2^, Paolo Gratton^2^, Renske E. Onstein^1^, Ammie K. Kalan^2^, Maureen S. McCarthy^2^, Gaëlle Bocksberger^2^, Lauren C. White^2^, Linda Vigilant^2^, Paula Dieguez^2^, Barrie Abdulai^3^, Thierry Aebischer^4^, Anthony Agbor^5^, Alfred Kwabena Assumang^6^, Emma Bailey^2^, Mattia Bessone^2,7^, Bartelijntje Buys^8^, Joana Silva Carvalho^7^, Rebecca Chancellor^9^, Heather Cohen^2^, Emmanuel Danquah^6^, Tobias Deschner^2^, Zacharie Nzooh Dongmo^10^, Osiris A. Doumbé^11^, Jef Dupain^12^, Chris S. Duvall^13^, Manasseh Eno-Nku^10^, Gilles Etoga^10^, Anh Galat-Luong^14^, Rosa Garriga^15^, Sylvain Gatti^16^, Andrea Ghiurghi^17^, Annemarie Goedmakers^6^, Anne-Céline Granjon^2^, Dismas Hakizimana^18^, Nadia Haydar^19^, Josephine Head^2^, Daniela Hedwig^20^, Ilka Herbinger^21^, Veerle Hermans^22,23^, Sorrel Jones^2^, Jessica Junker^1,2^, Parag Kadam^24^, Mohamed Kambi^2^, Ivonne Kienast^2^, Célestin Yao Kouakou^25^, Kouamé Paul N’Goran^10^, Kevin E. Langergraber^26,27^, Juan Lapuente^2,28^, Anne Laudisoit^29,30^, Kevin C. Lee^2,26^, Fiona Maisels^31,32^, Deborah Moore^†^, Bethan Morgan^32,33,34^, David Morgan^35^, Emily Neil^2^, Sonia Nicholl^2^, Louis Nkembi^36^, Anne Ntongho^10^, Christopher Orbell^37^, Lucy Jayne Ormsby^2^, Liliana Pacheco^38^, Alex K. Piel^7^, Lilian Pintea^39^, Andrew J. Plumptre^40^, Aaron Rundus^41^, Crickette Sanz^42,43^, Volker Sommer^44,45^, Tenekwetche Sop^1,2^, Fiona A. Stewart^7^, Jacqueline Sunderland-Groves^46^, Nikki Tagg^23^, Angelique Todd^47^, Els Ton^8^, Joost van Schijndel^8^, Hilde VanLeeuwe^31^, Elleni Vendras^2^, Adam Welsh^2^, José Francisco Carminatti Wenceslau^8^, Erin G. Wessling^48^, Jacob Willie^23^, Roman M. Wittig^2,22^, Nakashima Yoshihiro^49^, Yisa Ginath Yuh^2,50^, Kyle Yurkiw^2,51^ & Christophe Boesch^2^, Mimi Arandjelovic^2^, Hjalmar Kühl^1,2^

*^1^ German Centre for Integrative Biodiversity Research (iDiv), Halle-Jena-Leipzig, Deutscher Platz 5e, Leipzig 04103, Germany*

*^2^ Max Planck Institute for Evolutionary Anthropology, Department of Primatology, Deutscher Platz 6, Leipzig 04103, Germany*

*^3^ Energy Sector Utility Reform Project, 17E Wilkinson Road Freetown, Sierra Leone*

*^4^ Conservation et Plan d'aménagement de l’Aire de Conservation de Chinko, African Parks Network, Chinko Project, Kocho, RCA and active collaborator of the University of Fribourg, WegmannLab, Switzerland*

*^5^ African Parks Centurion Building, The Oval Corner Meadowbrook Lane and Sloane Street P.O. Box 2336, Lonehill 2062, RSA*

*^6^ Department of Wildlife and Range Management, Faculty of Renewable Natural Resources, Kwame Nkrumah University of Science and Technology, Kumasi, Ghana*

*^7^ School of Biological and Environmental Sciences, Liverpool John Moores University, Liverpool, UK*

*^8^ Chimbo Foundation, Huningspaed 6, 8567 LL Oudemirdum, Netherlands.*

*^9^ West Chester University, Depts of Anthropology & Sociology and Psychology, West Chester, PA, 19382 USA*

*^10^ World Wide Fund for Nature, Panda House Bastos, BP 6776 Yaounde, Cameroon*

*^11^ Sekakoh Organization, Bafoussam, Cameroon*

*^12^ Antwerp Zoo Foundation, Antwerp Zoo Society, Koningin Astridplein 20-26, 2018 Antwerpen, Belgium*

*^13^ Department of Geography and Environmental Studies, University of New Mexico, Albuquerque, NM 87131, USA*

*^14^ IRD (The French National Research Institute for Development), France*

*^15^ Tacugama Chimpanzee Sanctuary, PO Box 469, Freetown, Sierra Leone, West Africa*

*^16^ West African Primate Conservation Action (WAPCA), PO Box MB239, Accra GA-161-7942, Ghana*

*^17^ Independent Researcher, Via Passarelli 67, 00128 Rome, Italy*

*^18^ Department of Biology, University of Burundi, Burundi*

*^19^ Jane Goodall Institute Spain and Senegal, Dindefelo Biological Station, Dindefelo, Kedougou, Senegal*

*^20^ Elephant Listening Project, Center for Conservation Bioacoustics, Cornell Lab of Ornithology, Cornell University, 159 Sapsucker Woods Road, 14850 Ithaca, USA*

*^21^ WWF Germany, Reinhardtstr. 18, 10117 Berlin, Germany*

*^22^ Taï Chimpanzee Project, CSRS, BP 1301, Abidjan 01, Côte d’Ivoire*

*^23^ Centre for Research and Conservation, Antwerp Zoo Society, Koningin Astridplein 20-26, 2018 Antwerpen, Belgium*

*^24^ University of Cambridge, Pembroke Street, Cambridge, CB2 3QG, UK*

*^25^ Université Jean Lorougnon Guédé, BP 150 Daloa, Côte d’Ivoire*

*^26^ School of Human Evolution and Social Change, Arizona State University, Tempe, AZ, 85281, USA*

*^27^ Institute of Human Origins, Arizona State University, TEmpe, AZ, 85281, USA*

*^28^ Comoé Chimpanzee Conservation Project, Comoé National Park, Kakpin, Côte d’Ivoire*

*^29^ Ecohealth Alliance, 460 west 34th street, Ste1701, 10001 New York, USA*

*^30^ University of Antwerp, Campus Drie Eiken, lokaal D.133, Universiteitsplein 1 - 2610 Antwerpen, Belgium*

*^31^ Wildlife Conservation Society (WCS), Bronx, New York, USA*

*^32^ Faculty of Natural Sciences, University of Stirling, Stirling FK9 4LA, Scotland, UK*

*^33^ San Diego Zoo Global, 15600 San Pasqual Valley Road, Escondido, CA 92027-7000, USA*

*^34^ Ebo Forest Research Project, BP 3055, messa, Yaounde, Cameroon*

*^35^ Lester E Fisher Center for the Study and Conservation of Apes, Lincoln Park Zoo, Chicago, IL, 60614, USA*

*^36^ Environment and Rural Development Foundation, PO Box 189 Buea, Cameroon*

*^37^ Panthera, 8 W 40TH ST, New York, NY 10018, USA*

*^38^ WARA Conservation Project – GALF, République de Guinée*

*^39^ The Jane Goodall Institute, 1595 Spring Hill Road, Suite 550, Vienna, VA, 22182, USA*

*^40^ Key Biodiversity Area Secretariat, c/o BirdLife International, David Attenborough Building, Pembroke Street, Cambridge CB2 3QZ, UK*

*^41^ West Chester University, Department of Psychology, West Chester, PA, 19382 USA*

*^42^ Washington University in St. Louis, Department of Anthropology, Saint Louis, MO, 63130, USA*

*^43^ Wildlife Conservation Society, Congo Program, Brazzaville, Republic of Congo*

*^44^ Gashaka Primate Project, 663001 Serti, Taraba State, Nigeria*

*^45^ University College London, Department of Anthropology, London WC1E 6BT, UK*

*^46^ Faculty of Forestry, University of British Columbia, 2424 Main Mall, Vancouver, BC V6T 1Z4, Canada*

*^47^ WWF-CAR, Bangui, Central African Republic*

*^48^ Department of Human Evolutionary Biology, Harvard University, 11 Divinity Ave, 02138 Cambridge, MA, USA*

*^49^ College of Bioresource Science, Nihon University, 1866 Kameino, Fujisawa, Kanagawa 252-0880, Japan*

*^50^ University of Concordia, 1455 Boulevard de Maisonneuve O, Montréal, QC H3G 1M8, Canada*

*^51^ Pan Verus Project Outamba-Kilimi National Park, Sierra Leone*

*† Deceased*

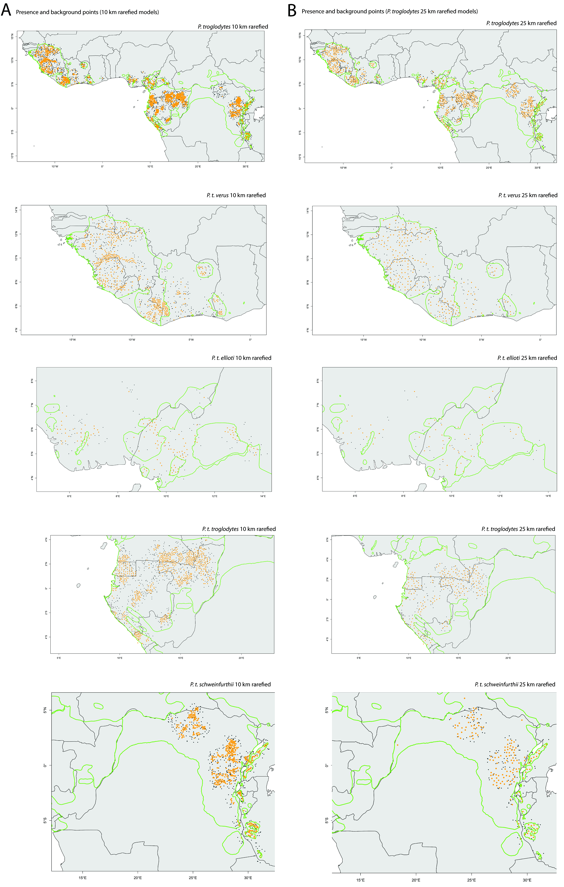

**Fig. S1.** Background points used to construct ensemble species distribution models for full species and each subspecies separately. Orange dots represent presence points, black dots are background points (randomly generated from within a 0.5 decimal degree radius of presence points). Country borders (dark grey lines) and chimpanzee subspecies ranges (green lines, Humle et al., 2018) also shown. A) Based on the 10 km rarefied spatial data as presences for species distribution models, B) Based on the 25 km rarefied spatial data as presences for species distribution models.

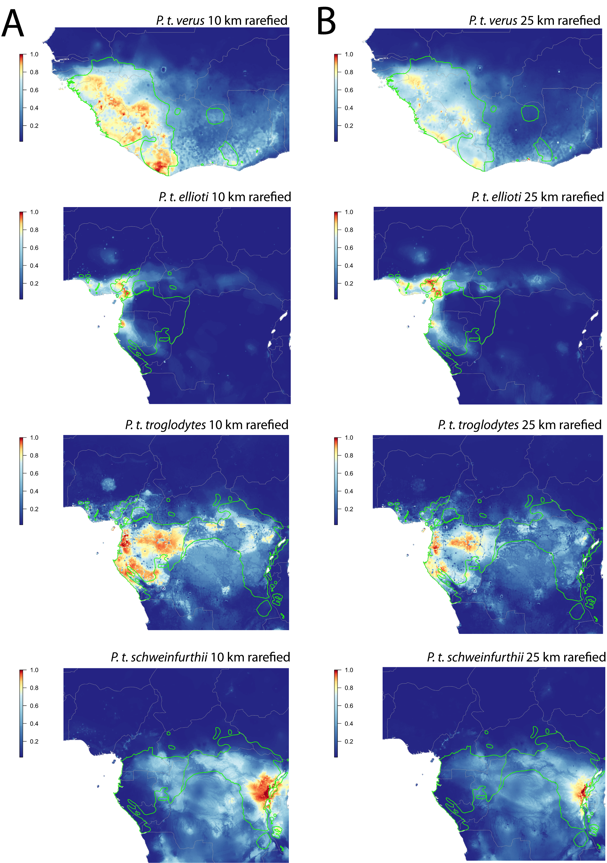

**Fig. S2.** Contemporary species distribution models for chimpanzees (0= low suitability to 1=highest suitability) using our final ensemble models. Country borders (grey lines) and chimpanzee subspecies ranges (green lines, Humle et al., 2018) also shown. A) Based on the 10 km rarefied spatial data as presences for species distribution models, B) Based on the 25 km rarefied spatial data as presences for species distribution models.

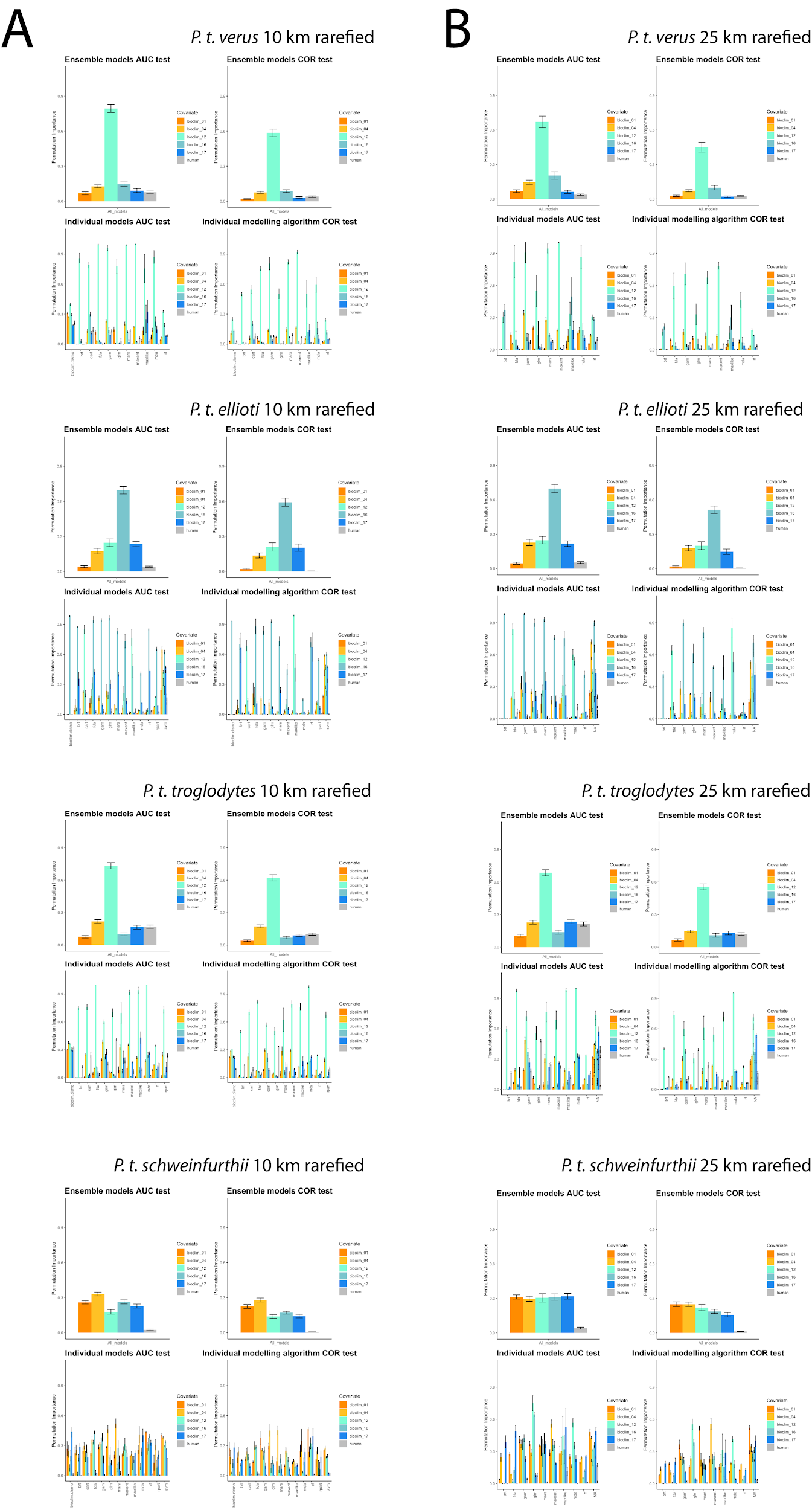

**Fig. S3.** Predictor variable permutation importance for chimpanzee contemporary species distribution models. Upper panels of each quartet show histograms of variable importance averaged across modelling algorithms, lower panels show variable importance per modelling algorithm. Left panels of each quartet show AUC permutation importance, Right panels show COR permutation importance. Axes represent variable importance (y axis), per modelling algorithm (x axis), bioclim_01 = mean annual temperature, bioclim_04 = temperature seasonality, bioclim_12 = mean annual precipitation, bioclim_16 = precipitation of the wettest quarter, bioclim_17 = precipitation of the driest quarter, human = human density. A) Based on the 10 km rarefied presence points, B) Based on the 25km rarefied presence points. for species distribution models.
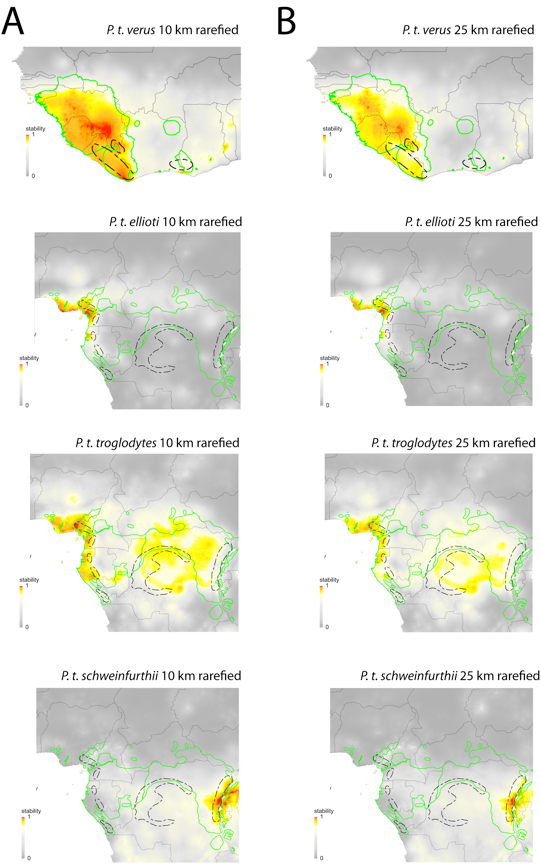

**Fig. S4.** Stability of chimpanzee habitat suitability (0= low suitability stability to 1=high suitability stability, for each subspecies modelled separately) over 62 snapshots of paleoclimate reconstructions representing the past 120,000 years using the Dynamic stability approach (dispersal rate of 5 m/year, amounting to a total possible dispersal of 600 km over the 120,000 years) compared to previous estimates of forest refugia (dotted black lines, Maley, 1996). A) Based on the 10 km rarefied presence points for species distribution models, B) Based on the 25 km rarefied presence points for species distribution models. Country borders (grey lines) and chimpanzee subspecies ranges (green lines, Humle et al., 2018) are also shown.

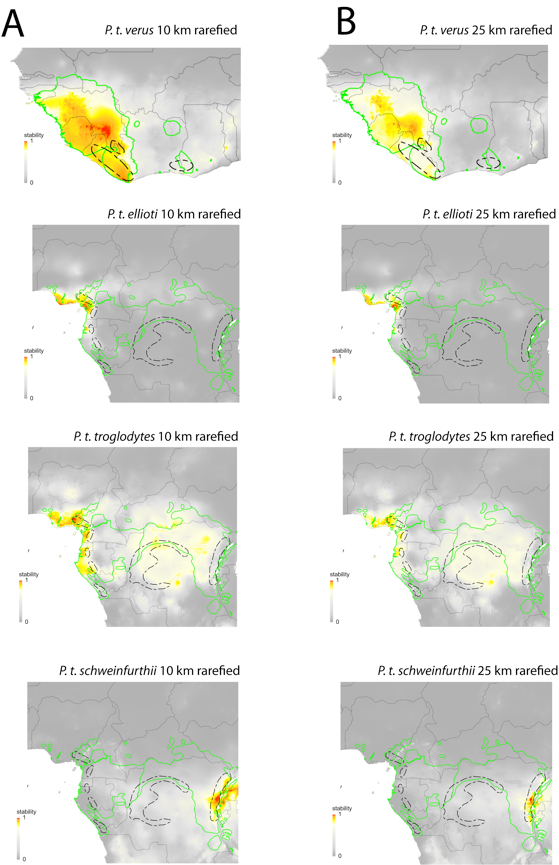

**Fig. S5.**  Stability of chimpanzee habitat suitability (0= low suitability stability to 1=high suitability stability, for each subspecies modelled separately) over 62 snapshots of paleoclimate reconstructions representing the past 120,000 years using the Static stability approach compared to previous estimates of forest refugia (dotted black lines, Maley, 1996). A) Based on the 10 km rarefied presence points for species distribution models, B) Based on the 25 km rarefied presence points for species distribution models. Country borders (grey lines) and chimpanzee subspecies ranges (green lines, Humle et al., 2018) are also shown.

**Table S1**. Performance of each individual modelling algorithm based on five replicates for full species (*P. troglodytes*) and each of the four currently recognised subspecies using presence points rarefied to minimum 25 km distance from one another. Numbers indicate Area Under the Curve of a Receiver Operating Characteristics plot (AUC) and True Skill Statistic (TSS) for each modelling algorithm. ***Bold italic*** values represent AUC < 0.8 or TSS < 0.5, the minimum threshold we set for that modelling algorithm to be included in the ensemble models. Modelling algorithm abbreviations – bioclim: Bioclim, bioclim.dismo: Bioclim from the dismo R package, brt: Boosted Regression Trees, cart: Classification and Regression Trees, fda: Flexible Discriminant Analysis, gam: Generalized Additive Model, glm, Generalized Linear Model, mars: Multivariate Adaptive Regression Spline, maxent: Maximum Entropy, maxlike: Maximum Entropy-like, mda: Mixture Discriminant Analysis, rf: Random Forest, rpart: Recursive Partitioning and Regression Trees, svm, Support Vector Machine.

| modelling algorithm | *P. troglodytes* |  | *P. t. verus* |  | *P. t. ellioti* |  | *P. t. troglodytes* | | *P. t. schweinfurthii* | |
| --- | --- | --- | --- | --- | --- | --- | --- | --- | --- | --- |
| evaluation metric | AUC | TSS | AUC | TSS | AUC | TSS | AUC | TSS | AUC | TSS |
| bioclim | ***0.59*** | **0.17** | ***0.61*** | **0.3** | 0.8 | **0.49** | ***0.72*** | **0.41** | ***0.7*** | **0.36** |
| bioclim.dismo | ***0.68*** | **0.33** | ***0.73*** | **0.44** | 0.84 | 0.66 | 0.87 | 0.72 | 0.8 | 0.58 |
| brt | 0.81 | 0.51 | 0.83 | 0.59 | 0.95 | 0.88 | 0.87 | 0.65 | 0.88 | 0.6 |
| cart | ***0.77*** | 0.56 | ***0.74*** | 0.59 | 0.83 | 0.69 | 0.86 | 0.69 | ***0.78*** | 0.59 |
| fda | 0.81 | 0.52 | 0.82 | 0.55 | 0.97 | 0.87 | 0.83 | 0.65 | 0.88 | 0.65 |
| gam | 0.86 | 0.61 | 0.85 | 0.61 | 0.91 | 0.81 | 0.94 | 0.8 | 0.91 | 0.71 |
| glm | 0.81 | 0.5 | 0.8 | 0.54 | 0.97 | 0.86 | 0.87 | 0.65 | 0.89 | 0.63 |
| mars | 0.86 | 0.59 | 0.84 | 0.61 | 0.96 | 0.9 | 0.93 | 0.76 | 0.9 | 0.67 |
| maxent | 0.85 | 0.58 | 0.86 | 0.61 | 0.97 | 0.89 | 0.94 | 0.79 | 0.92 | 0.71 |
| maxlike | 0.81 | 0.54 | 0.82 | 0.56 | 0.97 | 0.88 | 0.84 | 0.68 | 0.88 | 0.66 |
| mda | 0.82 | 0.53 | 0.8 | 0.51 | 0.98 | 0.91 | 0.87 | 0.69 | 0.86 | 0.64 |
| rf | 0.89 | 0.68 | 0.85 | 0.61 | 0.96 | 0.91 | 0.95 | 0.83 | 0.95 | 0.78 |
| rpart | ***0.72*** | 0.57 | ***0.78*** | 0.61 | 0.88 | 0.76 | 0.91 | 0.73 | ***0.79*** | 0.62 |
| svm | ***0.76*** | 0.63 | ***0.59*** | 0.59 | ***0.78*** | 0.84 | 0.94 | 0.79 | 0.87 | 0.64 |
| Total n models in ensemble | 9 |  | 9 |  | 13 |  | 13 |  | 11 |  |

**Table S2**. Pearson’s correlation coefficients between contemporary habitat suitability models, static stability and dynamic stability for full species (*P. troglodytes*) and each of the four currently recognised subspecies (10 km rarefied presence points vs. 25 km rarefied presence points).

|  | Contemporary habitat suitability | Static stability | Dynamic stability |
| --- | --- | --- | --- |
| *P. troglodytes* (full species) | 0.967 | 0.977 | 0.983 |
| *P. t. verus* | 0.975 | 0.976 | 0.98 |
| *P. t. ellioti* | 0.971 | 0.98 | 0.986 |
| *P. t. troglodytes* | 0.976 | 0.985 | 0.99 |
| *P. t. schweinfurthii* | 0.975 | 0.98 | 0.989 |
